# SERCA is a host target of the SARS-CoV-2 envelope protein linking calcium homeostasis to autophagy

**DOI:** 10.64898/2026.08.14.744854

**Authors:** Blanka Berta, Sarolta Tóth, Péter Lőrincz, Zurab Darjania, Nicolas Alexander Toranosuke Kato, Alaa Benachour, Nour Benachour, Tamás Hegedűs, Rita Padányi

## Abstract

The SARS-CoV-2 envelope (E) protein is a virulence factor that remodels host endomembranes, but mechanisms remain incompletely understood. We recently demonstrated that E protein interacts with and inhibits the sarco/endoplasmic reticulum Ca²⁺-ATPase (SERCA), disrupting ER calcium homeostasis. Here, we investigated how this perturbation affects autophagy-associated membrane organization. E protein expression induced lipidated LC3 accumulation and enlarged p62-positive structures, consistent with dysregulated autophagic turnover. Although E protein partially colocalized with LC3 and p62, enlarged p62-positive structures were also observed in cells retaining the reticular ER distribution of E protein, indicating that their formation does not require association with E protein or ER reorganization. E protein also increased the association of p62-positive structures with lysosomes without altering lysosome abundance. Pharmacological SERCA activation attenuated E protein-induced remodeling of autophagy-associated structures, demonstrating that SERCA inhibition contributes to these alterations. Together, our findings establish SERCA-dependent ER calcium homeostasis as a host pathway linking E protein expression to remodeling of autophagy-associated membrane compartments, providing a mechanistic framework for how the SARS-CoV-2 E protein promotes ER membrane remodeling associated with coronavirus replication.

## INTRODUCTION

Positive-strand RNA viruses extensively remodel host endomembranes to establish replication organelles that support viral genome replication while shielding viral intermediates from host defense mechanisms (Roingeard *et al*, 2022; Münz *et al*, 2025). In coronaviruses, these replication organelles primarily consist of endoplasmic reticulum (ER)-derived double-membrane vesicles (DMVs), whose formation depends on coordinated remodeling of ER membranes and host lipid homeostasis (Zimmermann *et al*, 2023; Roingeard *et al*, 2022; V’kovski *et al*, 2021). Although viral non-structural proteins such as nsp3 and nsp4 initiate ER membrane zippering, curvature and formation of the characteristic DMV-spanning pore (Zimmermann *et al*, 2023), increasing evidence indicates that multiple host pathways involved in ER organization, phospholipid homeostasis, and autophagy are essential for DMV biogenesis (Ji *et al*, 2022; Schneider *et al*, 2021). Genome-wide CRISPR screens identified TMEM41B and VMP1 as essential pan-coronavirus host factors (Schneider *et al*, 2021), and subsequent mechanistic studies demonstrated that these ER-resident proteins regulate phospholipid distribution and membrane remodeling required for efficient DMV formation (Ji *et al*, 2022; Tábara *et al*, 2018). Consequently, coronavirus replication is increasingly viewed as a process that depends on coordinated viral and host mechanisms governing ER membrane homeostasis (Münz *et al*, 2025).

Autophagy has emerged as one of the major cellular pathways manipulated during coronavirus infection (Münz *et al*, 2025). While autophagy normally contributes to cellular homeostasis through lysosomal degradation of cytoplasmic material, coronaviruses exploit selected components of the autophagy machinery while preventing complete degradative maturation (Khan *et al*, 2024). This uncoupling enables the virus to utilize autophagy-associated membrane remodeling while avoiding lysosomal degradation (Münz *et al*, 2025). Several viral proteins have been implicated in this process. The accessory protein ORF8 suppresses ER-phagy by recruiting ER-phagy receptors into p62 condensates, thereby promoting DMV formation (Tan *et al*, 2023), whereas expression of the envelope (E) protein induces LC3 lipidation, translational shutoff and autophagy-associated stress responses (Waisner *et al*, 2023). NSP6 has also been shown to promote early autophagosome formation while restricting autophagosome expansion and autophagosome–lysosome fusion, further supporting replication organelle biogenesis (Zhang *et al*, 2024, 2025; Cottam *et al*, 2014). More recently, naturally occurring mutations in E protein have been shown to alter viral susceptibility to autophagy, further strengthening the functional connection between E protein and autophagy regulation (Klute *et al*, 2025). Together, these findings suggest that distinct coronavirus proteins converge on autophagy-associated membrane remodeling through complementary mechanisms to facilitate viral replication.

SERCA occupies a central position within this regulatory network. Beyond maintaining ER calcium stores, SERCA coordinates ER calcium homeostasis with membrane contact dynamics, ER membrane organization and autophagic maturation (Tábara & Escalante, 2016; Tábara *et al*, 2018). VMP1 promotes SERCA activity to regulate ER membrane contact sites required for autophagosome biogenesis, placing SERCA upstream of both ER membrane remodeling and autophagy (Zhao *et al*, 2017; Zack *et al*, 2023). Its activity is modulated by endogenous regulins, including phospholamban (PLN), and disruption of SERCA regulation has been associated with defective autophagic maturation in diverse physiological settings (Vafiadaki *et al*, 2024). More recently, studies of PLN-R14del cardiomyopathy have suggested that disturbed SERCA regulation contributes not only to calcium dysregulation but also to abnormal sarco/endoplasmic reticulum (S/ER) organization, supporting a broader role of SERCA in maintaining ER membrane architecture (Vafiadaki *et al*, 2024). Notably, influenza A virus suppresses SERCA activity, contributing to impaired autophagic maturation, whereas pharmacological activation of SERCA restores autophagic flux and attenuates virus-induced cellular dysfunction (Peng *et al*, 2021).

The coronavirus envelope (E) protein is a small ER-localized transmembrane protein that functions as an essential virulence factor (Liao *et al*, 2006). Besides its well-characterized viroporin activity, E protein predominantly localizes to the ER-Golgi intermediate compartment (ERGIC), where coronavirus assembly and replication organelle formation occur, placing it in an ideal position to influence ER membrane homeostasis (Liao *et al*, 2006; Waisner *et al*, 2023; Schoeman & Fielding, 2019). We recently demonstrated that SARS-CoV-2 E protein directly binds SERCA2b and inhibits SERCA-mediated ER Ca²⁺ uptake by acting as a viral exoregulin that competes with endogenous SERCA regulators (Berta *et al*, 2024). These findings raised the possibility that E protein-mediated SERCA inhibition influences not only ER calcium homeostasis but also the broader membrane remodeling pathways coordinated by SERCA.

Here, we investigated whether direct inhibition of SERCA by SARS-CoV-2 E protein perturbs the SERCA-dependent ER membrane regulatory network and thereby contributes to autophagy-associated membrane remodeling. We show that E protein induces accumulation of LC3-II and p62 together with enlarged p62-positive structures, and that pharmacological activation of SERCA partially rescues these alterations. Our findings support a model in which SARS-CoV-2 E protein perturbs a SERCA-dependent regulatory network linking ER calcium homeostasis, membrane organization and autophagic maturation, providing a mechanistic framework that connects SERCA inhibition with ER membrane remodeling during coronavirus infection.

## RESULTS

### E protein dysregulates autophagic turnover

Because SERCA-dependent ER Ca²⁺ homeostasis contributes to autophagy regulation, we examined whether SARS-CoV-2 E protein expression alters autophagy in human lung-derived epithelial A549 cells. Autophagy was monitored by immunoblotting for LC3 and p62/SQSTM1. LC3-II represents the lipidated, autophagic membrane-associated form of LC3, generated from cytosolic LC3-I upon autophagy induction. The p62 abundance provides a complementary readout of autophagic cargo turnover, as it is degraded together with autophagic cargo during the autophagic process. Starvation and chloroquine served as reference conditions for autophagy induction and impaired autophagic degradation, respectively (Klionsky *et al*, 2008).

Expression of E protein markedly increased LC3-II levels, reaching values comparable to those induced by starvation (Figure 1A, C). In contrast, p62 exhibited a strikingly different pattern. Whereas starvation reduced p62 levels, as expected, E protein expression resulted in pronounced p62 accumulation (Figure 1A, B). Notably, starvation failed to reduce this E protein-induced accumulation of p62, indicating that E protein overrides the normal autophagic response to nutrient deprivation. Thus, although E protein increased LC3-II to levels comparable to those observed during starvation, it elicited a distinct autophagic response, marked by the accumulation rather than depletion of p62.

**Figure 1.**
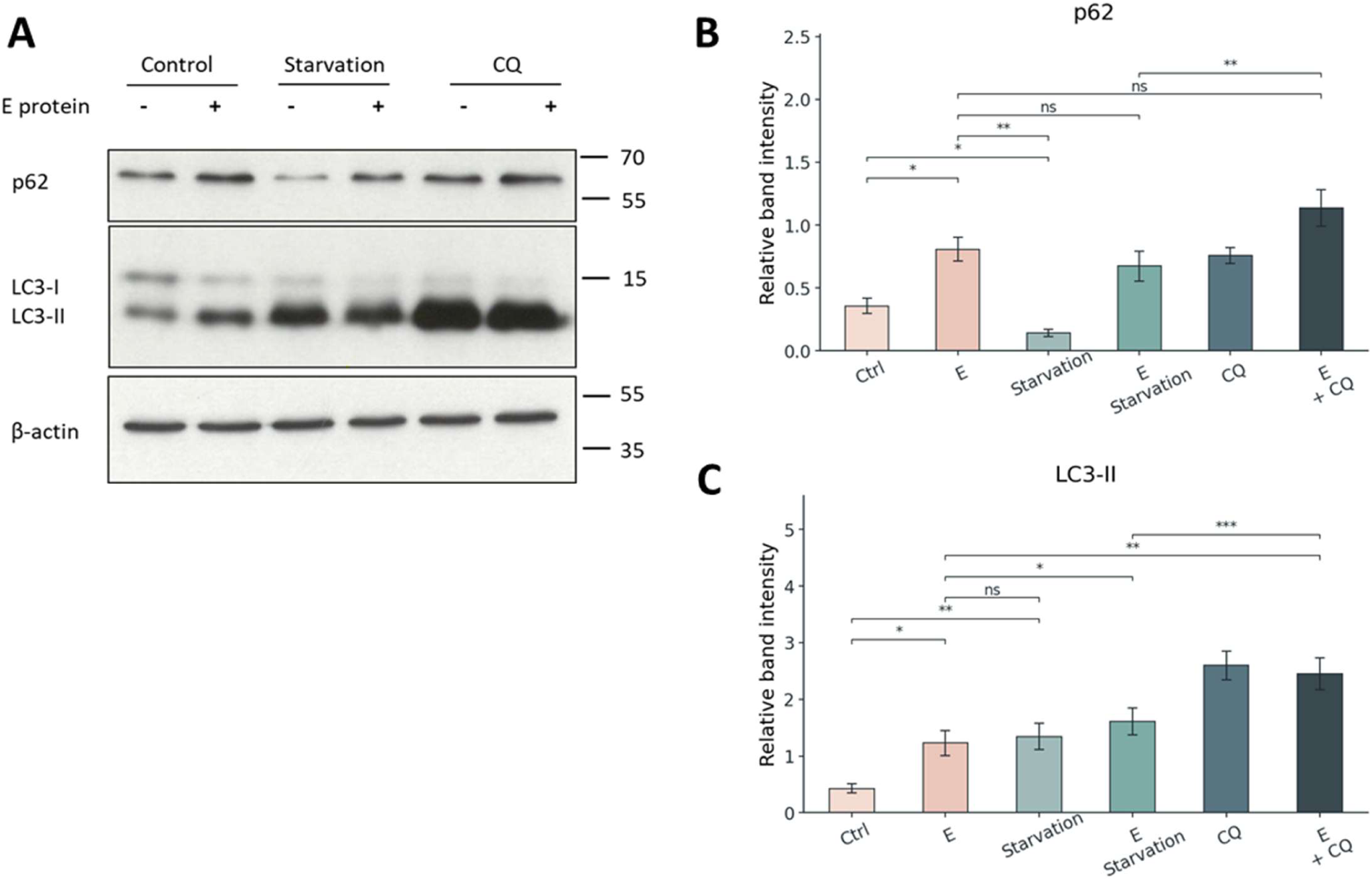
SARS-CoV-2 E protein induces an abnormal autophagy-related phenotype in A549 cells. **(A)** Representative Western blot showing p62/SQSTM1 and LC3 in control and SARS-CoV-2 E protein-expressing A549 cells under basal conditions, starvation, and starvation plus chloroquine (CQ) treatment. β-actin served as a loading control. **(B)** Quantification of p62/SQSTM1 protein levels normalized to β-actin. **(C)** Quantification of LC3-II protein levels normalized to β-actin. Data are presented as mean ± SEM from 4–5 independent experiments. Statistical significance was determined using paired two-tailed Student’s *t* tests. *p < 0.05; **p < 0.01; ns, not significant.

To determine whether autolysosomal degradation was completely blocked, cells were treated with chloroquine. Chloroquine further increased both LC3-II and p62 levels in E protein-expressing cells (Figure 1A–C), indicating that lysosome-dependent turnover of these autophagy markers remained active in E protein-expressing cells, suggesting that autophagic flux was partially impaired rather than completely blocked. Consistent with these findings, analysis of the LC3-II/LC3-I ratio showed a relative shift toward the LC3-II form in E protein-expressing cells, both under starvation and chloroquine treatment, although the differences did not reach statistical significance, with substantial variability observed between experiments (Figure S1A).

Together, these findings demonstrate that E protein induces the accumulation of autophagy-related structures while partially impairing their clearance.

### E protein induces the accumulation of autophagosome-like structures

To determine whether the biochemical changes were accompanied by morphological alterations, autophagy markers were examined by confocal microscopy. In control cells, LC3 and p62 staining was predominantly diffuse, with only occasional punctate structures (Figure 2A). In contrast, E protein-expressing cells contained numerous LC3- and p62-positive puncta, in agreement with the Western blotting results. E protein partially colocalized with both LC3 and p62, indicating that a fraction of the protein is associated with these structures (Figure S2). Notably, p62-positive puncta were readily detected even in cells displaying the characteristic ER localization of E protein, indicating that their formation does not require obvious alterations in E protein distribution or ER morphology.

**Figure 2.**
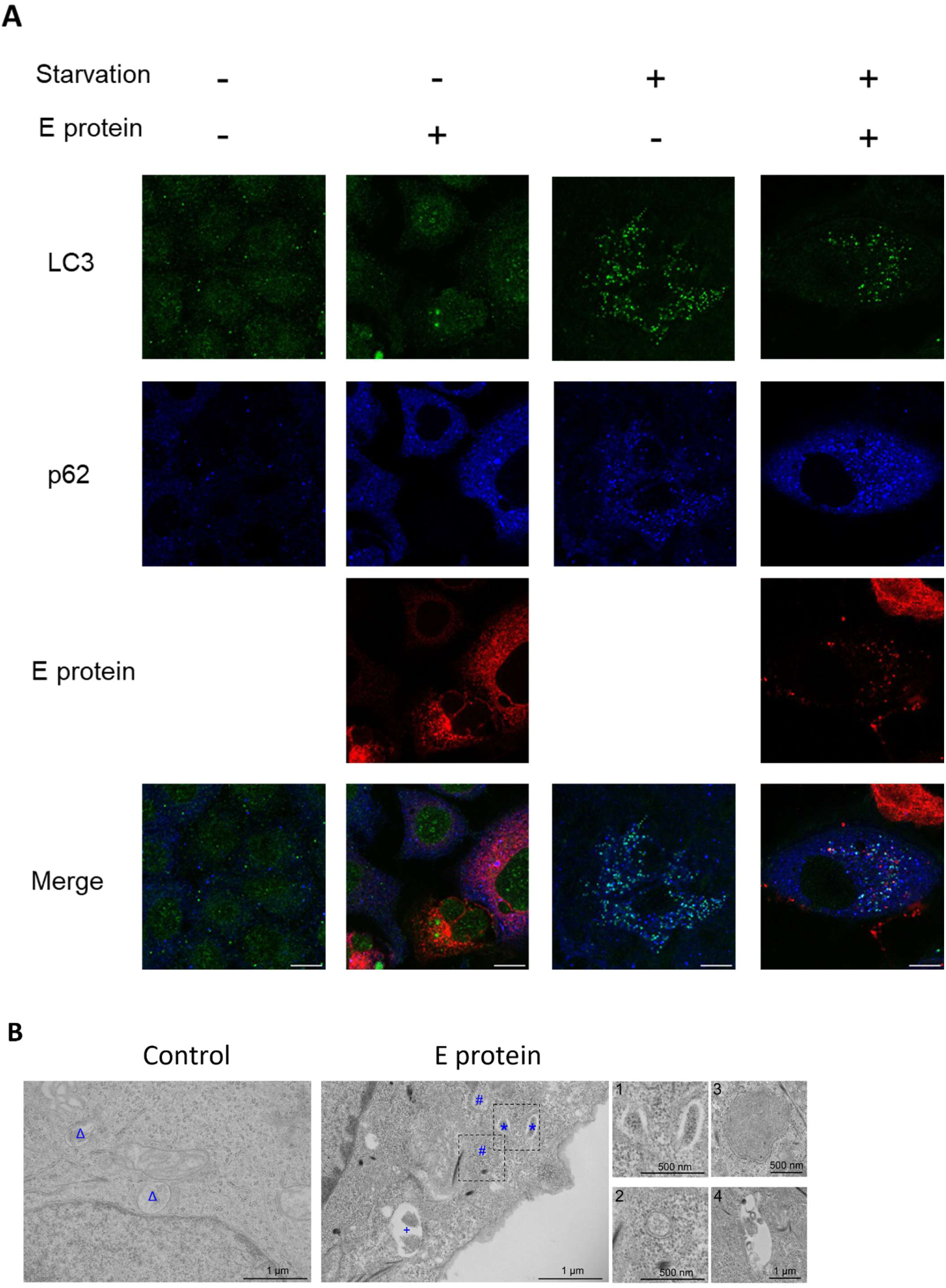
SARS-CoV-2 E protein induces the accumulation of autophagy-associated structures in A549 cells. **(A)** Representative confocal images of LC3, p62/SQSTM1, and SARS-CoV-2 E protein in control and E protein-expressing A549 cells under basal and starvation conditions. **(B)** Representative transmission electron micrographs of control and E protein-expressing A549 cells. Blue triangles indicate late endosomes/multivesicular bodies, asterisks indicate phagophores, plus signs indicate autophagosomes, and hash symbols indicate double-membrane vesicles (DMVs). Insets show higher-magnification views of the indicated structures. Scale bars: 10 μm in confocal images; values are indicated in the electron micrographs.

Transmission electron microscopy was used to further characterize the ultrastructural changes associated with E protein expression. Control A549 cells predominantly displayed normal-appearing vesicular structures, such as endosomal and lysosomal structures, whereas autophagic structures were rarely observed. In contrast, E protein-expressing cells contained multiple early autophagic structures, including phagophore-like membranes and autophagosomes, consistent with the accumulation of autophagy-related compartments observed by confocal microscopy. Double-membrane vesicular structures were also observed in E protein-expressing cells (Figure 2B) consistent with the extensive membrane remodeling associated with coronavirus proteins. In addition, enlarged autolysosome-like structures were frequently observed in E protein-expressing cells (Figure S3), consistent with altered lysosomal homeostasis and potentially impaired degradation. Together, these ultrastructural findings support the accumulation of autophagy-related and DMV structures and indicate altered lysosomal morphology.

Together, the confocal and ultrastructural analyses demonstrate that E protein promotes the accumulation of autophagosome-like structures in A549 cells.

### E protein inhibits SERCA-dependent ER Ca²⁺ refilling

We previously demonstrated that SARS-CoV-2 E protein inhibits SERCA activity in HeLa cells (Berta *et al*, 2024). To determine whether this effect is preserved in pulmonary cells, we examined SERCA-dependent ER Ca²⁺ refilling in A549 cells following ER store depletion with the ER-localized fluorescent Ca²⁺ indicator ER-GCaMP6-150.

Our experiments show that E protein expression significantly reduced the rate of ER Ca²⁺ refilling compared with control cells (Figure 3A, B), indicating impaired SERCA-dependent Ca²⁺ uptake. In contrast, starvation modestly but significantly increased the ER Ca²⁺ refilling rate compared with control cells, consistent with previous reports linking autophagy induction to enhanced SERCA activity (Zack *et al*, 2023). This finding suggests that autophagy induction itself does not impair ER Ca²⁺ refilling and therefore is unlikely to account for the reduced refilling observed in E protein-expressing cells. To determine whether the reduced ER Ca²⁺ refilling resulted from SERCA inhibition, E protein-expressing cells were treated with the SERCA activator CDN1163, which we expect to counterbalance the E protein inhibition. Indeed, CDN1163 significantly increased the ER Ca²⁺ refilling rate, restoring it to control levels (Figure 3B).

**Figure 3.**
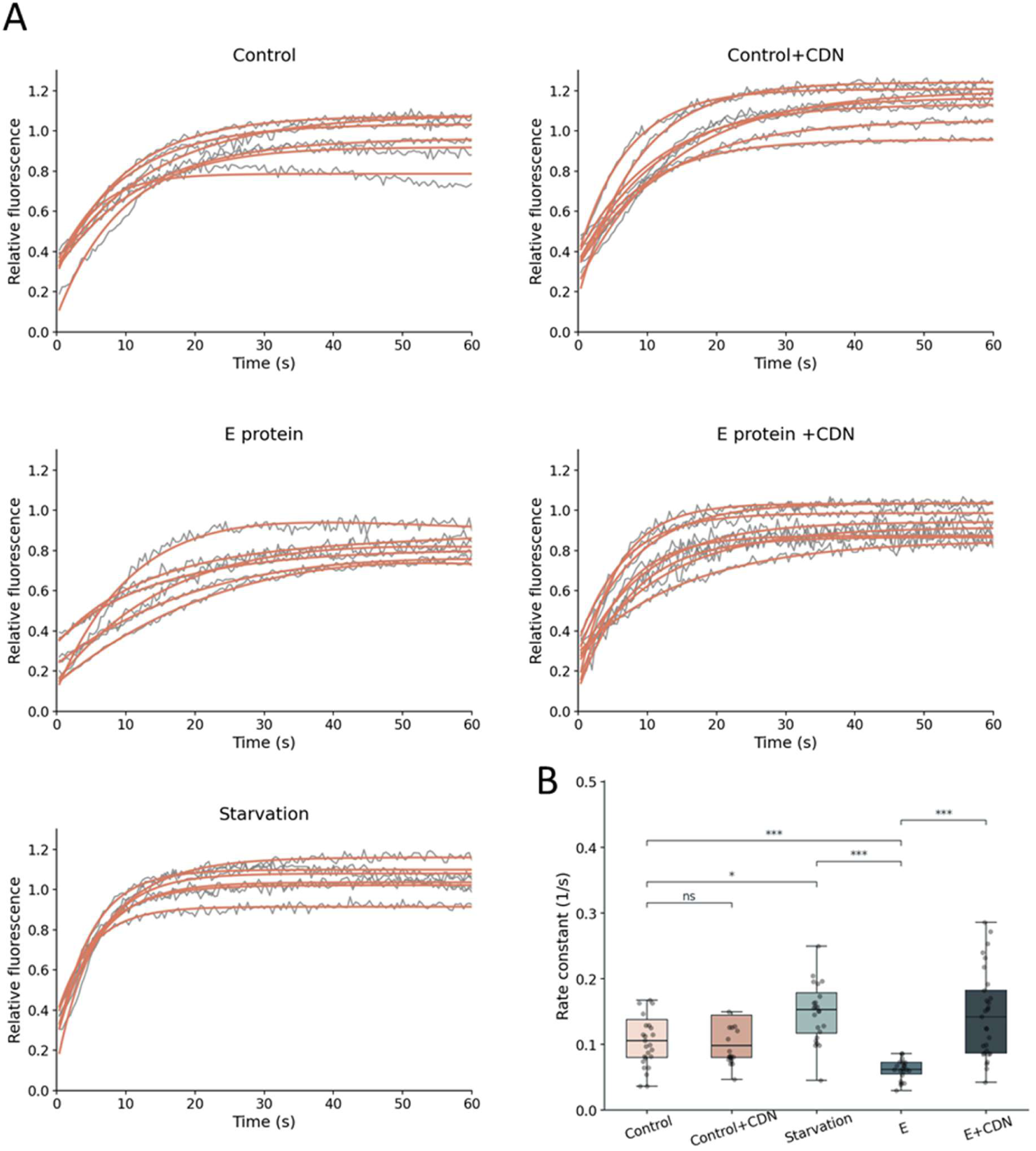
SARS-CoV-2 E protein inhibits SERCA-dependent ER Ca²⁺ refilling in A549 cells. **(A)** Representative ER Ca²⁺ refilling traces recorded after ATP-induced ER Ca²⁺ release and extracellular Ca²⁺ re-addition in control, starved, and SARS-CoV-2 E protein-expressing A549 cells, with or without CDN1163 treatment. **(B)** Quantification of ER Ca²⁺ refilling rate constants. Boxplots show the median and interquartile range; whiskers indicate 1.5 × IQR. Individual points represent single-cell measurements. Statistical significance was determined using the Kruskal–Wallis test followed by planned pairwise Mann–Whitney *U* tests. Statistical significance is indicated on the graph as follows: *p < 0.05, **p < 0.01, ***p < 0.001; ns, not significant. Only planned pairwise comparisons are indicated on the graph.

These findings demonstrate that E protein inhibits SERCA-dependent ER Ca²⁺ refilling in A549 cells and that pharmacological SERCA activation effectively reverses this defect.

### Altered cytosolic Ca²⁺ signaling is consistent with SERCA inhibition

Reduced SERCA activity is expected to alter intracellular Ca²⁺ homeostasis beyond ER refilling. We therefore analyzed ATP-evoked cytosolic Ca²⁺ responses by measuring both the peak amplitude and the area under the curve (AUC) using a fluorescent Ca²⁺-indicator, R-GECO expressed in the cytosol. Representative ATP-evoked cytosolic Ca²⁺ traces are shown in Figure 4A.

**Figure 4.**
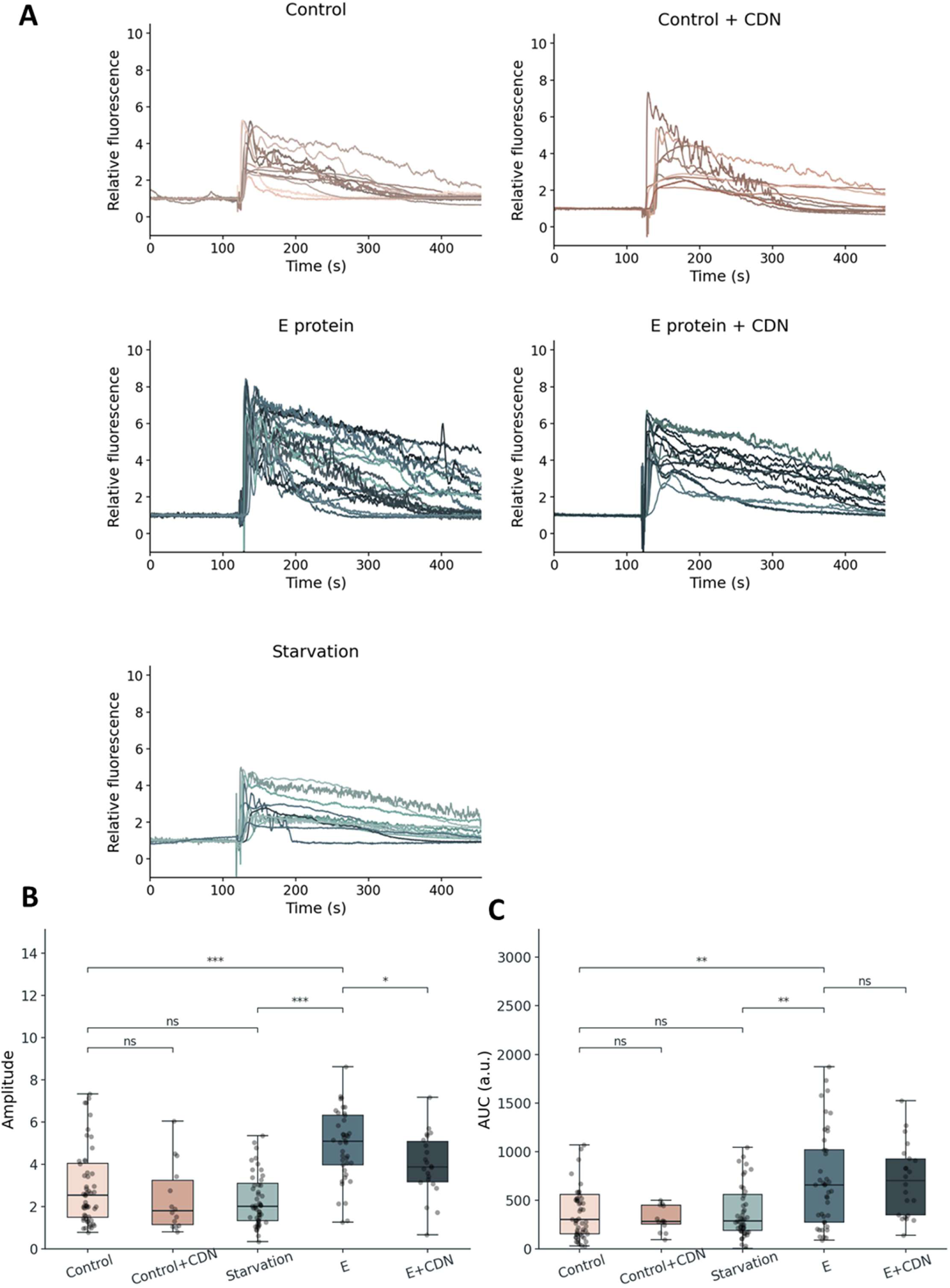
SARS-CoV-2 E protein alters ATP-evoked cytosolic Ca²⁺ signaling in A549 cells. **(A)** Representative ATP-evoked cytosolic Ca²⁺ traces recorded in control, starved, and SARS-CoV-2 E protein-expressing A549 cells, with or without CDN1163 treatment. **(B)** Quantification of the peak amplitude of ATP-induced cytosolic Ca²⁺ responses. **(C)** Quantification of the area under the curve (AUC) of ATP-induced cytosolic Ca²⁺ responses. Boxplots show the median and interquartile range; whiskers indicate 1.5 × IQR. Individual points represent single-cell measurements. Statistical significance was determined using the Kruskal–Wallis test followed by planned pairwise Mann–Whitney *U* tests. Only planned pairwise comparisons are indicated on the graph. Statistical significance is indicated as follows: *p < 0.05, **p < 0.01, ***p < 0.001; ns, not significant.

E protein expression significantly increased both the peak amplitude and the AUC of ATP-induced cytosolic Ca²⁺ responses compared with control cells (Figure 4A–C). In contrast, starvation did not significantly affect either parameter, consistent with the ER Ca²⁺ refilling measurements.

Treatment with the SERCA activator CDN1163 significantly reduced the peak amplitude of ATP-induced Ca²⁺ responses in E protein-expressing cells, whereas the reduction in AUC did not reach statistical significance (Figure 4B, C), indicating only partial restoration of cytosolic Ca²⁺ signaling.

Together, the ER refilling and cytosolic Ca²⁺ measurements support the conclusion that E protein disrupts cellular Ca²⁺ homeostasis through SERCA inhibition.

### SERCA activation partially rescues the E protein-induced autophagy phenotype

To determine whether impaired SERCA-dependent ER Ca²⁺ refilling contributes to the autophagy phenotype induced by E protein expression, we pharmacologically restored SERCA activity using the SERCA activator CDN1163 and assessed the resulting autophagy phenotype. Cells were treated with CDN1163 for 48 h following transfection, after which autophagy markers were analyzed by Western blotting. CDN1163 treatment did not significantly alter the E protein-induced increase in LC3-II levels (Figure 5A, B), indicating that restoration of SERCA activity had little effect on LC3-II accumulation under these experimental conditions. Consistently, CDN1163 did not significantly alter the LC3-II/LC3-I ratio in E protein-expressing cells (Figure S1B).

**Figure 5.**
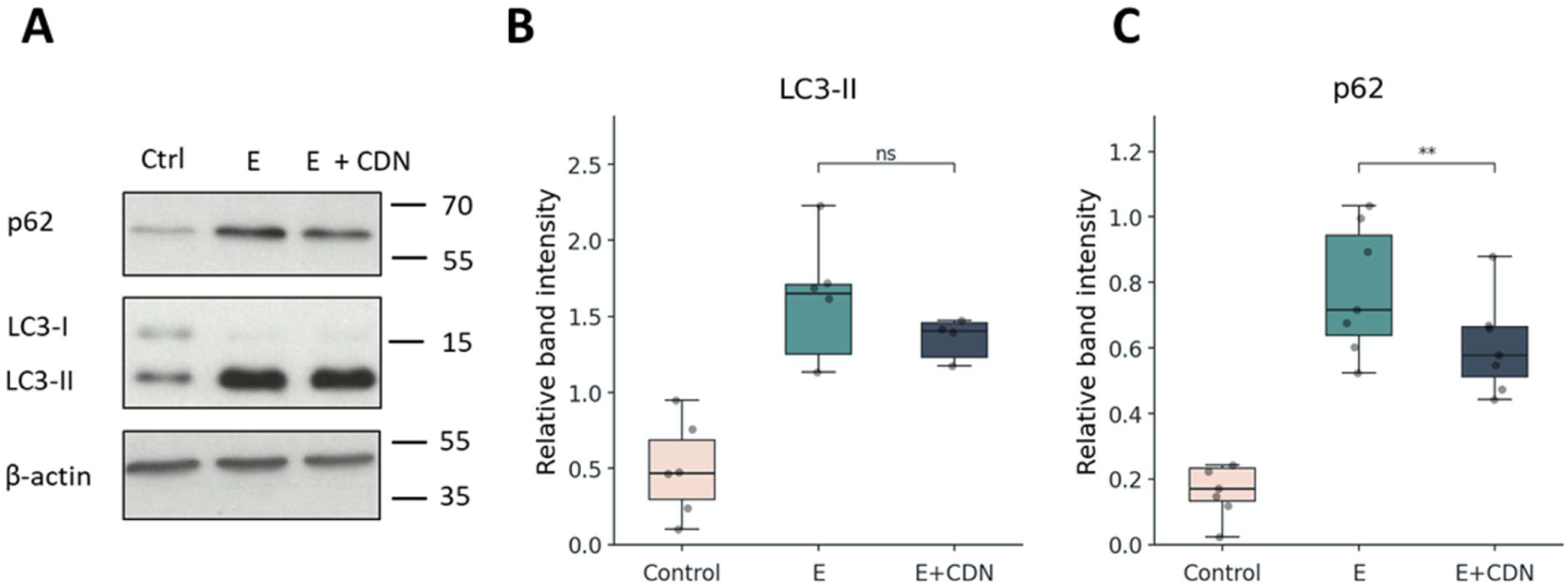
SERCA activation partially reduces E protein-induced p62 accumulation. **(A)** Representative Western blot showing LC3, p62/SQSTM1 and β-actin in control, SARS-CoV-2 E protein-expressing, and CDN1163-treated E protein-expressing A549 cells. **(B)** Quantification of LC3-II protein levels normalized to β-actin. **(C)** Quantification of p62/SQSTM1 protein levels normalized to β-actin. Boxplots show the median and interquartile range; individual points represent independent experiments. Statistical significance was determined using paired two-tailed Student’s *t* tests. *p < 0.05, **p < 0.01; ns, not significant.

In contrast, CDN1163 significantly reduced p62 levels in E protein-expressing cells (Figure 5A, C), although p62 remained elevated compared with control cells. This observation demonstrates that SERCA activation partially rescued the E protein-induced autophagy phenotype.

Together, these results indicate that SERCA inhibition contributes to the E protein-induced accumulation of p62 but is not solely responsible for the overall autophagy phenotype.

Confocal microscopy was also used to confirm these biochemical findings. E protein expression induced the accumulation of enlarged p62-positive structures, whereas treatment with CDN1163 visibly reduced their abundance (Figure 6A). Analysis of p62 puncta size distribution revealed that, compared with starvation, which was characterized by predominantly small p62-positive puncta, E protein expression shifted the distribution toward larger structures. CDN1163 preferentially reduced the abundance of the largest p62-positive compartments (Figure 6B). To quantify this effect, only p62-positive structures larger than 0.75 μm² were included in the analysis. CDN1163 significantly reduced the abundance of these enlarged p62-positive structures (Figure 6C).

**Figure 6.**
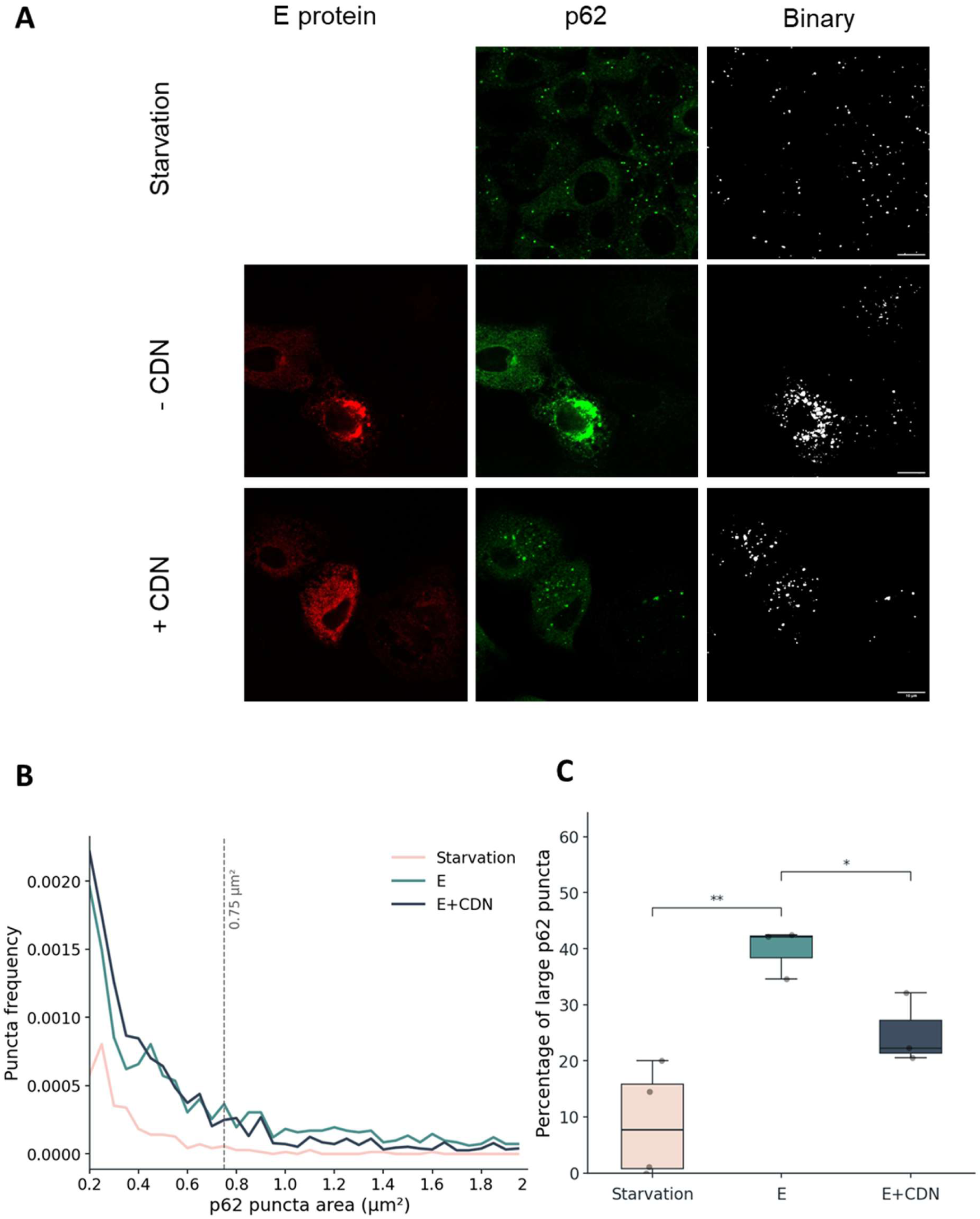
SERCA activation reduces the accumulation of enlarged p62-positive structures. **(A)** Representative confocal images of p62/SQSTM1 in starved control and SARS-CoV-2 E protein-expressing A549 cells with or without CDN1163 treatment. Binary masks show the segmented p62-positive objects used for quantitative analysis. **(B)** Size distribution of p62-positive structures. The dashed line indicates the 0.75 µm² threshold used to define enlarged p62-positive structures. **(C)** Quantification of enlarged p62-positive structures (>0.75 µm²) expressed as a percentage of all p62-positive puncta ≥0.2 µm². Statistical significance was determined using planned Welch’s *t* tests with Holm correction for multiple comparisons. Statistical significance is indicated as follows: *p < 0.05, **p < 0.01; ns, not significant. Scale bars: 10 μm.

Together, these findings indicate that SERCA activation partially limits the accumulation of enlarged p62-positive structures induced by E protein.

### E protein enhances the association of p62-positive structures with lysosomes

To determine whether the accumulated p62-positive structures progress to the lysosomal stage of the autophagy pathway, lysosomes were visualized by immunostaining for LAMP1 (Figure 7A). Representative Western blot analysis did not reveal an apparent difference in LAMP1 protein levels between control and E protein-expressing cells (Figure S4D). Consistent with this, the number of LAMP1-positive vesicles did not differ significantly between the groups (Figure 7B), indicating that E protein does not increase overall lysosome abundance. Pearson’s correlation analysis revealed a significant increase in the spatial association between p62 and LAMP1 signals in E protein-expressing cells (Figure S4B). However, because this test is influenced by signal intensity and object morphology, and because LAMP1-positive lysosomes frequently surrounded rather than directly overlapped with p62-positive structures, colocalization was further evaluated using an object-based approach (Bolte & Cordelières, 2006; Dunn *et al*, 2011). Object-based colocalization analysis likewise demonstrated a significant increase in the percentage of LAMP1-positive objects associated with p62-positive structures (Figure 7C), confirming that the accumulated p62-positive compartments are closely associated with lysosomes. Complementary object-based analyses of the reciprocal p62–LAMP1 association and nearest-neighbor distance are shown in Figure S4A, C.

**Figure 7.**
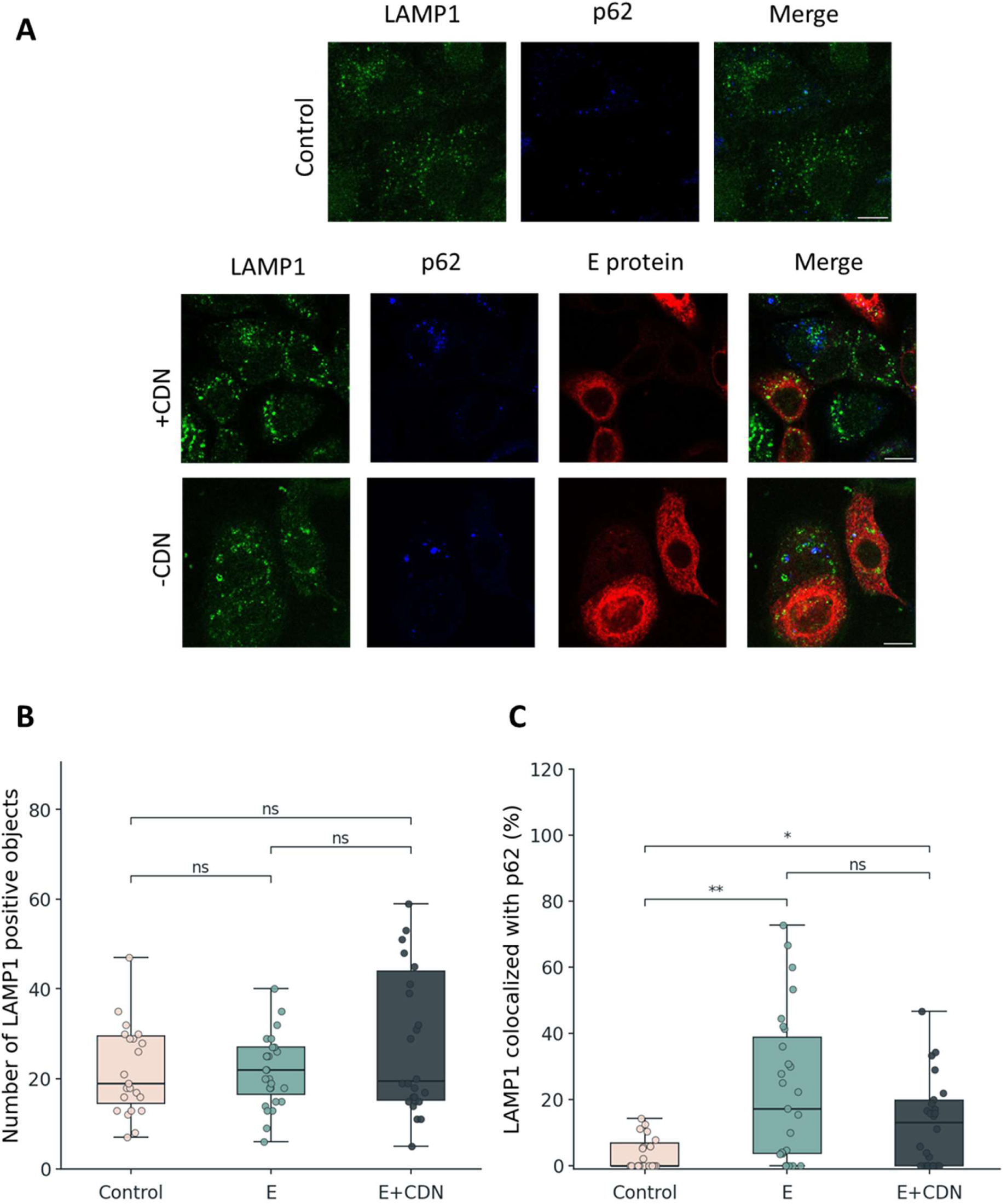
E protein increases the association of LAMP1-positive structures with p62-positive compartments, which is not significantly altered by SERCA activation. **(A)** Representative confocal images of LAMP1, p62/SQSTM1, and SARS-CoV-2 E protein in control and E protein-expressing A549 cells with or without CDN1163 treatment. **(B)** Quantification of LAMP1-positive objects per image. **(C)** Object-based colocalization analysis showing the percentage of LAMP1-positive objects associated with p62-positive structures. Boxplots show the median and interquartile range; whiskers indicate 1.5 × IQR. Individual points represent images. Statistical significance was determined using the Kruskal–Wallis test followed by planned pairwise Mann–Whitney *U* tests. Only planned pairwise comparisons are indicated on the graph. Significance values shown on the graphs are based on uncorrected p values: **p < 0.01; ns, not significant. Scale bars: 10 μm.

Importantly, treatment with CDN1163 did not significantly affect either the number of LAMP1-positive vesicles or the object-based association between p62-positive structures and lysosomes (Figure 7B, C). These findings indicate that SERCA inhibition contributes to the early accumulation of abnormal autophagy-related structures, whereas the enhanced association of these structures with lysosomes is mediated through an additional SERCA-independent mechanism.

## DISCUSSION

The SARS-CoV-2 envelope (E) protein is an essential virulence factor that perturbs ER homeostasis, yet the molecular pathways linking E protein function to autophagy-associated membrane remodeling have not been clearly defined. We previously identified SERCA2b as a direct molecular target of E protein by demonstrating their interaction and showing that E protein inhibits SERCA activity (Berta *et al*, 2024). Here, we extend these findings by demonstrating impaired SERCA function in pulmonary epithelial cells and showing that the consequences of E protein-mediated SERCA inhibition extend beyond altered ER Ca²⁺ handling to autophagy-associated compartments. E protein expression is accompanied by marked accumulation of p62-positive structures together with increased LC3-II levels, consistent with altered autophagic processing. Importantly, pharmacological restoration of SERCA activity significantly reduced p62 accumulation, indicating that SERCA-dependent ER calcium homeostasis contributes to the regulation of autophagic cargo processing. Together, these findings identify impaired SERCA activity as a mechanistic link between E protein-induced ER dysfunction and altered autophagic maturation.

The concurrent increase in LC3-II and p62 observed by immunoblotting suggests that autophagy-associated structures accumulate without being efficiently cleared (Mauvezin *et al*, 2015). This interpretation is further supported by the presence of enlarged p62-positive structures, which likely reflect altered organization of p62-positive autophagy-associated compartments rather than increased p62 synthesis alone. Although activation of SERCA partially alleviated this phenotype, it did not fully restore normal p62 levels, indicating that SERCA inhibition represents an important, but not exclusive mechanism through which E protein remodels the endomembrane system. Additional activities of E protein, including its viroporin function, induction of ER stress and interactions with host membrane proteins, are also likely to contribute to the observed phenotype (Liao *et al*, 2006; Shaban *et al*, 2021). Consistent with this interpretation, E protein expression did not alter overall lysosome abundance, yet increased the association of p62-positive structures with LAMP1-positive compartments. These observations argue against a defect in lysosome availability and instead suggest that autophagic cargo reaches LAMP1-positive compartments, while its subsequent lysosomal processing or clearance may be altered. The enlarged autolysosome-like structures observed in E protein-expressing cells by electron microscopy (Supplementary Figure S3) are consistent with altered lysosomal homeostasis, although this morphological observation alone does not establish a defect in lysosomal degradation (Lee *et al*, 2010; Festa *et al*, 2018; Chandrasekaran *et al*, 2021).

Our observation that E protein affects both ER Ca²⁺ handling and the reorganization of autophagy-associated compartments suggests that its consequences extend beyond a single Ca²⁺ transport process. Consistent with broader effects of E protein on organelle organization, SARS-CoV-2 E protein expression has also been reported to alter ER–mitochondria contact sites and mitochondrial Ca²⁺ handling (Poggio *et al*, 2023). VMP1, an ER-resident protein required for autophagosome biogenesis, promotes SERCA activity and regulates ER membrane contact sites that are essential for autophagic maturation (Zack *et al*, 2023). Consistent with this concept and our results, studies have identified VMP1 and TMEM41B as essential host factors for coronavirus replication, where they regulate ER membrane organization and double-membrane vesicle (DMV) biogenesis (Ji *et al*, 2022). VMP1-dependent activation of SERCA has been proposed to promote the proper detachment of isolation membranes from the ER during autophagosome formation, linking SERCA activity directly to ER membrane remodeling (Zhao *et al*, 2017). Together, these observations suggest that SERCA-dependent ER calcium homeostasis is integrated into a broader regulatory network controlling ER membrane organization, membrane dynamics and autophagic maturation (Kohler *et al*, 2020). Our findings place SARS-CoV-2 E protein within this emerging framework by suggesting that direct inhibition of SERCA perturbs this interconnected regulatory network.

This interpretation is further supported by independent studies in unrelated biological systems. The cardiomyopathy-associated phospholamban mutant PLN-R14del, which chronically inhibits SERCA, causes p62 and LC3 accumulation through defective autophagosome–lysosome fusion (Vafiadaki *et al*, 2024). During influenza A virus infection, the viral M2 protein, a small oligomeric protein that, similarly to SARS-CoV-2 E protein, disrupts intracellular ion homeostasis, contributes to defective autophagic degradation, which is accompanied by impaired SERCA activity. Pharmacological activation of SERCA partially restored autophagic maturation and attenuated virus-induced cellular dysfunction (Peng *et al*, 2021; Shaban *et al*, 2021). Although these systems differ fundamentally from SARS-CoV-2 infection, the convergence of pharmacological, genetic and viral evidence suggests that disturbed SERCA regulation represents a common upstream determinant of defective autophagic maturation.

Despite lacking obvious sequence homology, influenza A virus M2 and SARS-CoV-2 E protein are both small oligomeric viroporins that localize to intracellular membranes, perturb host ion homeostasis and contribute to viral pathogenicity (Pinto *et al*, 1992; Liao *et al*, 2006; Cao *et al*, 2021). Whereas influenza A virus M2 has been functionally linked to SERCA inhibition, SARS-CoV-2 E protein has been shown to interact directly with SERCA2b and to behave as a viral exoregulin by occupying an interaction interface overlapping with endogenous SERCA regulins (Berta *et al*, 2024). This observation raises the possibility that viral membrane proteins may exploit conserved host regulatory mechanisms in addition to functioning as ion channels. Rather than acting solely by altering membrane permeability, E protein appears to modulate an endogenous regulatory pathway controlling ER calcium homeostasis and membrane organization.

Our findings therefore support a broader interpretation of the relationship between SERCA and autophagy. Rather than identifying another viral regulator of autophagy, our findings place SERCA-dependent ER calcium homeostasis upstream of the autophagic remodeling induced by the SARS-CoV-2 E protein. Within this framework, altered autophagic maturation, p62 condensate formation and ER membrane remodeling represent interconnected consequences of disturbed ER homeostasis. Because ER membrane remodeling is essential for coronavirus replication organelle biogenesis (Zimmermann *et al*, 2023; Roingeard *et al*, 2022), perturbation of SERCA-dependent ER homeostasis may contribute to creating a membrane environment favorable for viral replication.

Although the present study used ectopic E protein expression rather than viral infection, this controlled system allowed E protein to be introduced independently of other viral components and enabled its specific effects on SERCA function and autophagy-associated compartments to be examined. At the same time, ectopic expression may not fully reflect the temporal regulation, relative abundance, and broader cellular context of E protein during authentic infection. Therefore, the contribution of E protein-mediated SERCA inhibition to coronavirus replication remains to be established. In addition, further ultrastructural analyses will be neededto determine whether the enlarged p62-positive structures are directly associated with membrane intermediates involved in replication organelle formation.

In conclusion, our findings identify impaired SERCA activity as a mechanistic link between SARS-CoV-2 E protein expression and altered autophagic maturation. Together with accumulating evidence implicating SERCA, VMP1 and TMEM41B in ER membrane organization (Schneider *et al*, 2021; Ji *et al*, 2022), our results support an emerging model in which disruption of SERCA-dependent ER calcium homeostasis contributes to the membrane remodeling processes exploited by coronaviruses. This work extends our previous identification of E protein as a viral SERCA regulator by demonstrating that its functional consequences extend beyond ER calcium handling to include remodeling of autophagy-associated membrane compartments.

## METHODS

### Cell culture and transient transfection

A549 cells (CCL-185, ATCC) were maintained in Dulbecco’s modified Eagle’s medium (DMEM) supplemented with 10% fetal bovine serum and Penicillin/Streptomycin at 37 °C in a humidified atmosphere containing 5% CO₂.

For biochemical experiments, cells were seeded into 6-well plates at a density of 3 × 10⁵ cells per well in 2 ml medium one day prior to transfection. For immunofluorescence microscopy, cells were seeded into 8-well Ibidi chambers (Ibidi 80807) at a density of 3.5 × 10⁴ cells per well in 200 µl medium one day prior to transfection.

Cells were transiently transfected with the previously described N-terminally mCherry-tagged SARS-CoV-2 E protein construct (Berta *et al*, 2024) using Lipofectamine 3000 transfection reagent (Thermo Fisher Scientific, L3000008) according to the manufacturer’s instructions. Non-transfected cells processed in parallel under the same experimental conditions were used as controls. For immunofluorescence experiments, cells were fixed with 4% paraformaldehyde in PBS for 15 min at 37 °C.

### Starvation, chloroquine and CDN1163 treatments

For starvation-induced autophagy, cells were incubated in serum- and glutamine-free EBSS for 3 h before fixation or sample collection. Where indicated, chloroquine (CQ) was applied at a final concentration of 50 µM in serum- and glutamine-free EBSS for 3 h during starvation.

For SERCA activation experiments, cells were treated with 10 µM CDN1163 diluted from a 5 mM stock solution prepared in DMSO. CDN1163 treatment was started 2 h after transfection and maintained until fixation or sample collection. Untreated control and E protein-expressing cells were processed in parallel under the same experimental conditions.

### Western blot analysis

Western blotting was used to assess autophagy-related protein levels in control and SARS-CoV-2 E protein-expressing A549 cells following starvation, chloroquine treatment and/or SERCA activation. After the indicated treatments, cells were collected by TCA precipitation and processed for SDS-PAGE. Proteins were separated by electrophoresis on 15% SDS-polyacrylamide gels and transferred to polyvinylidene difluoride (PVDF) membranes.

Non-specific binding sites were blocked by incubating the membranes in blocking solution containing 5% non-fat dry milk in Tris-buffered saline with Tween-20 (TBST) for 1 h. Membranes were then incubated overnight at 4 °C with the appropriate primary antibodies. The following primary antibodies were used: anti-LC3A/B (Cell Signaling Technology, #12741S; 1:1000 in BSA-containing TBST), anti-p62/SQSTM1 (D5L7G, Cell Signaling Technology, #88588; 1:500 in BSA-containing TBST), and anti-β-actin (Sigma-Aldrich A1978; 1:20,000).

After washing in TBST, membranes were incubated with the corresponding HRP-conjugated secondary antibodies for 1 h at room temperature. HRP-conjugated anti-rabbit and anti-mouse secondary antibodies were used at 1:10,000 (Jackson ImmunoResearch 111-035-003 and 115-035-003). After washing, HRP-antibody-antigen complexes were detected using ECL.

LC3-II and p62/SQSTM1 band intensities were quantified and normalized to β-actin. Relative protein levels were calculated from independent experiments and are presented as indicated in the figure legends.

### Immunofluorescence staining and confocal microscopy

For immunofluorescence microscopy, A549 cells were grown in 8-well Ibidi chamber slides and treated as described above. After the indicated treatments, cells were fixed with 4% paraformaldehyde (PFA) for 15 min at 37°C. After washing with PBS, cells were permeabilized with methanol for 1 min and blocked for 1 h at room temperature in blocking solution containing 2 mg/ml BSA, 0.1% Triton X-100 and 5% donkey serum in PBS.

Cells were incubated with primary antibodies diluted in blocking solution for 1 h at room temperature. The following primary antibodies were used: anti-LC3A/B (Cell Signaling Technology, #12741S; 1:100), anti-p62/SQSTM1 (D5L7G, Cell Signaling Technology, #88588; 1:500), and anti-LAMP1/CD107a (Cell Signaling #9091; 1:100). After washing with PBS, cells were incubated with the appropriate fluorescent secondary antibodies diluted in blocking solution for 1 h at room temperature. Alexa Fluor 488-conjugated anti-rabbit secondary antibody was used at 1:500, and anti-mouse STAR RED secondary antibody was used at 1:250.

Fluorescence imaging was performed using a Nikon Eclipse Ti2 confocal microscope equipped with either a 20× objective or a 60× oil-immersion objective, depending on the experiment.

### Live-cell Ca²⁺ imaging and analysis

For live-cell Ca²⁺ imaging experiments, A549 cells were seeded into 8-well chambered cover glasses at a density of 0.6 × 10⁵ cells per well in 500 µl medium one day before transfection. Cells were transfected with either the cytosolic Ca²⁺ indicator R-GECO (Zhao *et al*, 2011) or the ER-targeted Ca²⁺ indicator ER-GCaMP6-150 (Juan-Sanz *et al*, 2017), with or without the mCherry-tagged SARS-CoV-2 E protein construct, using Lipofectamine 3000 as described above. Measurements were performed 24 h after transfection, following the indicated starvation or pharmacological treatments.

Live-cell fluorescence measurements were performed under temperature-controlled conditions using a Nikon Eclipse Ti2 confocal microscope equipped with a 20× objective.

For cytosolic Ca²⁺ measurements, cells expressing R-GECO were washed once with HBSS and maintained in 100 µl HBSS-based imaging buffer containing 2 mM CaCl₂ and 10 mM HEPES, pH 7.4. Following a 2-min baseline recording, ATP was added at a final concentration of 100 µM to evoke cytosolic Ca²⁺ signals. Cytosolic Ca²⁺ responses were quantified based on the amplitude and area under the curve.

For ER Ca²⁺ release and refilling measurements, cells expressing ER-GCaMP6-150 were washed and maintained in low-Ca²⁺ HBSS-based imaging buffer containing 10 mM HEPES, 100 µM CaCl2 and 100 µM EGTA, pH 7.4. Following a 2-min baseline recording, ATP was added at a final concentration of 100 µM to induce ER Ca²⁺ release. After ER store depletion, Ca²⁺-containing HBSS was added to restore the extracellular Ca²⁺ concentration to approximately 2 mM, and ER Ca²⁺ refilling was monitored.

Fluorescence intensities were extracted using the Time Series Analyzer plugin in ImageJ. Fluorescence traces were normalized to the mean baseline fluorescence (F₀) calculated from the initial 2-min recording period (F/F₀). Quantitative analyses were performed using custom Python scripts (Spyder IDE). For cytosolic Ca²⁺ measurements, response amplitude was calculated as the difference between the baseline and peak fluorescence, and the area under the curve (AUC) was calculated by integrating the normalized fluorescence signal over the first 6 min following ATP stimulation. For ER Ca²⁺ measurements, the refilling phase was fitted with a single-exponential function, and the corresponding rate constant was used to quantify SERCA-dependent ER Ca²⁺ refilling.

### Image analysis and quantification

Confocal images were analyzed using Fiji/ImageJ v1.54p (Schneider *et al*, 2012). For p62 size analysis, the area of individual p62-positive objects was measured. Objects smaller than 0.2 µm² were excluded, and enlarged p62-positive structures were defined as objects larger than 0.75 µm². Puncta-size distributions were normalized to the analyzed cell area, and the proportion of enlarged p62-positive objects was calculated relative to all detected objects larger than 0.2 µm².

LAMP1-positive object numbers and LAMP1–p62 colocalization were quantified using the JACoP plugin in Fiji/ImageJ (Bolte & Cordelières, 2006). Colocalization was assessed using Pearson’s correlation coefficient and object-based analysis. Object-based colocalization was expressed as the percentage of LAMP1-positive objects associated with p62-positive structures.

### Electron microscopy

Cells were fixed in 3.2% paraformaldehyde, 0.5% glutaraldehyde, 1% sucrose, and 2mM CaCl2 in 0.1 M sodium cacodylate, pH 7.4, overnight at 4°C, postfixed in 0.5% osmium tetroxide for 1 hour and in 2% aqueous uranyl acetate for 30 minutes. Samples were dehydrated in a graded series of ethanol and embedded in Durcupan ACM (Sigma 44611-44614) according to the manufacturer’s instructions. Ultrathin sections were stained in Reynold’s lead citrate and examined on a transmission electron microscope (JEM-1011; JEOL, Tokyo, Japan) equipped with a digital camera (Morada; Olympus) using iTEM software (Olympus) (Simon-Vecsei *et al*, 2021).

### Statistical analysis

Statistical analyses were performed using Spyder (Spyder version: 6.1.5 (standalone), Python version: 3.12.11 64-bit) with the SciPy (scipy.stats) package. Data are presented as mean ± SEM unless otherwise indicated. Pairwise comparisons were performed using two-tailed paired Student’s t tests, two-tailed Welch’s t tests, or Mann–Whitney U tests, as appropriate. Comparisons among multiple groups were performed using the Kruskal–Wallis test followed by planned pairwise Mann–Whitney U tests. The statistical test used for each experiment is indicated in the corresponding figure legend. Statistical significance was defined as P < 0.05.

## DATA AVAILABILITY

**Supplementary information:** The online version contains supplementary material.

## AUTHOR CONTRIBUTIONS

RP and TH conceived and developed the ideas. RP and BB designed and optimized the experiments. BB performed the majority of the experiments, with contributions from ZD, NK, AB and NB. LP and ST performed the electron microscopy experiments. ZD, NK, AB and NB also participated in data analysis. RP and BB analyzed and interpreted the data. RP and BB wrote the manuscript. RP, BB, TH and ST contributed to the critical reading and revision of the manuscript. TH acquired funding and supervised the project.

## ACKNOWLEDGEMENTS

We are grateful to Ágnes Enyedi for providing the R-GECO calcium sensor and the anti-LC3 antibody. We thank Krisztina Szendefyné Lór and I. Répássy for excellent technical assistance.

